# An improved epigenetic age estimation with TFMethyl Clock reveals DNA methylation changes during aging in transcription factor binding sites

**DOI:** 10.1101/2025.10.07.680024

**Authors:** Tushar Patel, Robert Schwarz, Konstantin Riege, Miri Varshavsky, Hans A. Kestler, Tommy Kaplan, Steve Hoffmann, Alena van Bömmel

## Abstract

Methylation-based epigenetic clocks are among the most accurate tools for predicting chronological age. Although DNA methylation (DNAm) at genomic CpG sites is linked to various regulatory mechanisms, the biological interpretability of epigenetic clocks remains surprisingly limited. One primary mechanism by which DNAm is thought to influence gene regulation is by modulating transcription factor binding activity. In this study, we examine established epigenetic clocks to assess the regulatory potential of their predictive CpGs during the aging process. Our analysis reveals that generally most CpG sites used by epigenetic clocks do not overlap known transcription factor binding sites (TFBSs), indicating that changes in TFBS dynamics may not account for prediction accuracy of these models. On the other hand, by identifying age-associated CpGs that overlap TFBSs, we identified transcription factors that may be involved in the aging process. Specifically, the TFBSs of *ZBED1, NFE2, CEBPB, FOXP1, EGR1, SP1, PAX5,* and *MAZ* were particularly enriched for age-associated CpGs, while *RBPJ, NFIC, RELA, IKZF1, STAT3,* and *USF2* were significantly protected against methylation changes. By focusing on TFBS-associated CpGs, combined with additional feature selection and engineering steps, we developed an alternative, *TFMethyl Clock* model, outperforming several existing approaches. Target genes of model-selected, age-predictive CpGs are enriched in the interleukin-1b production and long- chain fatty acid metabolism pathways. In contrast, these CpGs themselves are enriched mainly at binding sites of *NR2C2* TF. Furthermore, approximately three-fourths of the target genes downstream of age- predictive CpGs exhibit significant age-related changes, suggesting that our approach captures deeper insights into possible methylation-driven biological aging processes. Our findings demonstrate that incorporating regulatory loci into the design of epigenetic predictors may provide mechanistic insights into the aging process while maintaining or even improving the predictive power.

## Introduction

DNAm has been repeatedly shown to be a robust biomarker to predict chronological age [1];[2];[3]. While the first generation of epigenetic clocks [1];[2] already exhibited a remarkable accuracy, second- generation clocks incorporated further data to even obtain predictions on mortality or life-span [4];[5]. Fueled by these encouraging results, the palette of epigenetic clock models has become increasingly diverse in recent years, capturing different aspects of DNAm during aging [6];[7];[8];[9]. The ever-increasing availability of training samples will likely further enhance the models’ accuracy in predicting chronological age [10]. However, a striking limitation of virtually all clock models is their inability to provide deeper insights into the biological aging process. In general, finding potentially causal and verifiable downstream biological mechanisms associated with learned features remains a challenging task [3];[11];[12]. This is particularly critical if such models are applied to assess the effects of interventions aimed at improving longevity or healthspan [11].

So far, most DNAm clocks have been developed using regularized regression methods such as elastic net [13] or LASSO [14]. During the typical training phase on a specific sample cohort, a relatively small set of CpGs is identified, whose methylation levels, combined and weighted in a linear model, are enough to predict chronological age [15]. Similar to other problems addressed with machine learning (ML) strategies, training aging clocks involves a large number of features (p) but relatively few samples (n). Typically, aging clocks are trained on 10^4^ - 10^5^ features (often correlated among themselves) with only around 10^3^ samples. To address the *big p, little n* or “p>>n” problem, regularization terms can help achieve sufficient model performance with a minimal set of features by shrinking the weights of all other features to zero, effectively removing them from the equation. Depending on the chosen regularization method and its parameters, even small changes in methylation values in the training samples can lead to the selection of different sets of CpGs with little overlap [12].

Besides selection stability problems, features that don’t exhibit a particularly high correlation with chronological age may still be selected during training–a phenomenon referred to as noise accumulation [16];[17]. Consequently, the biological interpretability of epigenetic clocks may be limited. While the pre- selection of features is a frequently suggested strategy to mitigate instabilities and the accumulation of noisy features [16], it may also be suitable to focus on particular aspects of gene regulation, e.g., on the expression of transposable elements (TEs) [18]. Further, to deplete confounders and reduce prediction bias for specific age groups, the DNAm clocks should restrict the modelling on CpG sites exhibiting aging-associated DNAm signals [7];[19].

On the other hand, the mechanism by which DNAm is thought to influence gene regulation is through modifying transcription factor (TF) binding activity [20]. In particular, DNAm may sterically hinder or selectively enhance transcription factor (TF) binding at regulatory elements, depending on the specific properties of the TF [21]. Therefore, to improve the biological interpretability of the epigenetic clocks, it may be helpful to focus on CpGs overlapping known TF binding sites (TFBS).

Here, we present a framework for building a regulatory predictor, *TFMethyl Clock,* whose features provide a better insight into the molecular mechanisms of aging. The proposed feature selection step focuses on aging-correlated CpG sites located within experimentally known TFBSs. Furthermore, our modeling approach is preceded by a feature clustering step to enhance the robustness of predictions. Subsequently, we investigate the extent to which TFBSs are affected by age-specific methylation changes and characterize their potential target genes.

## Results

Predictive performance and biological interpretability of epigenetic clocks To scrutinize the biological underpinnings of established epigenetic clocks, we investigated their model CpGs in terms of their regulatory potential and direct aging correlations (Fig. 1A). Using DNAm data from the human blood aging compendium [22], we calculated the proportion of CpGs overlapping with experimentally validated TFBS (see Methods) and proportions of CpGs with a strong chronological age correlation (Rank-based Spearman’s measure |⍴| > 0.5).

**Figure 1.**
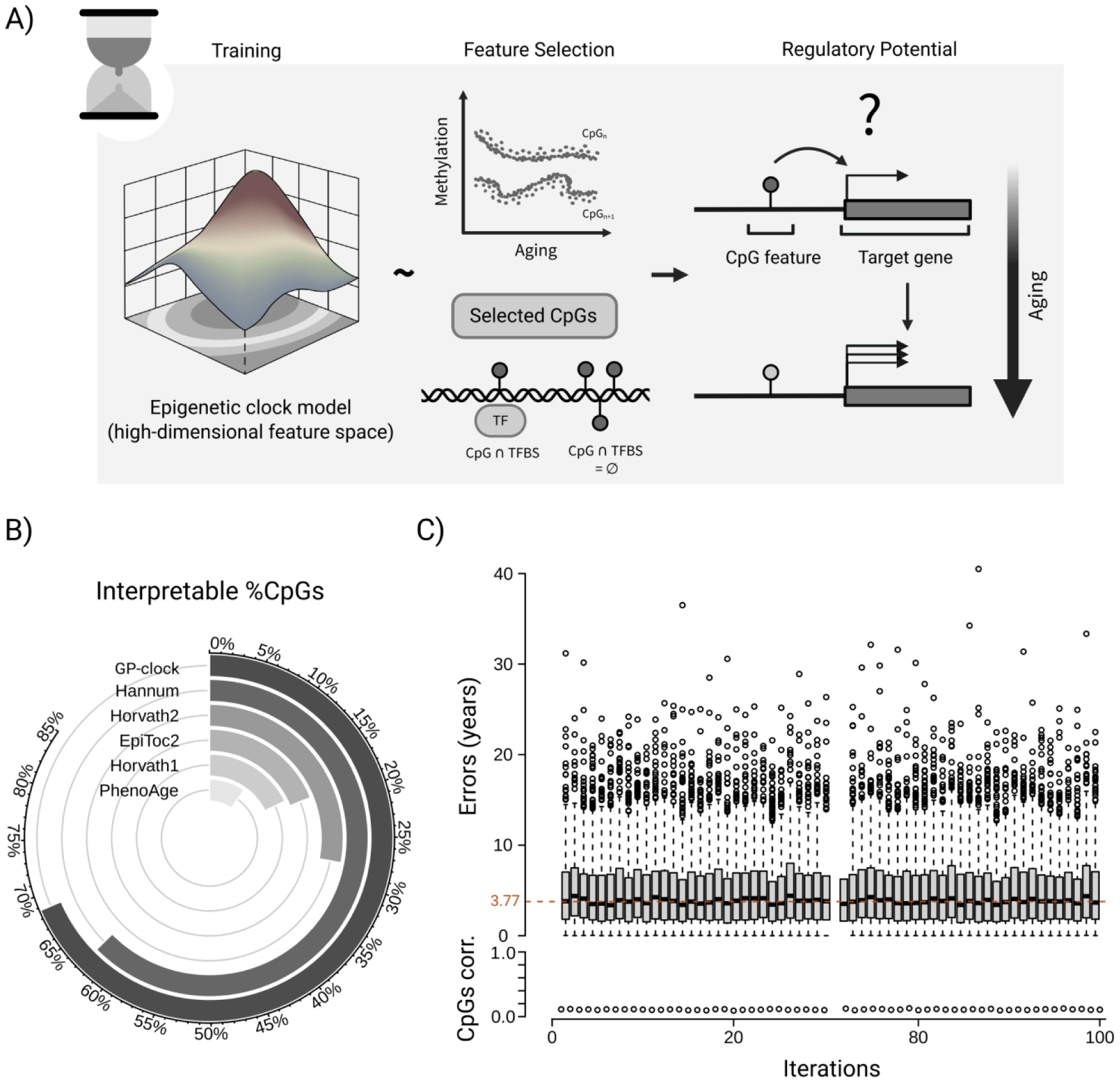
Systematic assessment of epigenetic clocks for regulatory potential and age prediction performance. A) Conceptual framework for dissecting epigenetic clock features by their biological impact. A trained epigenetic clock model is defined in a high-dimensional CpG feature space (left). A subset of model-selected CpG sites (middle) shows strong or limited aging signatures, and possibly TFBS at their loci (or both). Regulatory potential of selected CpGs for downstream gene regulation by correlating their DNAm change with target gene expression change during aging (right) B) Proportion of biologically interpretable CpGs in the clock models. The circular bar plot represents the percentage of aging-correlated (|⍴| > 0.5) and TFBS overlapping CpGs in the contemporary epigenetic clocks. C) Absolute errors of epigenetic clock models trained on biologically less informative features (CpGs with age |⍴| < 0.5 & non-overlapping TFBS) tested on the validation cohort GSE84727. Boxes represent the IQR of absolute errors, with the median of all models highlighted as a colored dashed line (MdAE = 3.77 yrs). Dot plot below shows the respective average age-correlation score for the CpGs used for model training.

For the analyzed methylation dataset, the Horvath clocks contain 18% (Horvath1) [2] and 28% (Horvath2) [23] model CpGs strongly correlated with age and overlapping with a TFBS at the same time (Fig. 1B). For the healthspan-optimized PhenoAge model [4], we find that only 10% of CpGs fulfill both criteria. In turn, with 69% (49 out of 71 CpGs), the Gaussian-process clock model (GP-clock) [22] shows the most substantial proportion of correlated and TFBS-associated CpGs. A closer look at age correlations reveals that about 70% of PhenoAge and 60% of Horvath1 CpGs correlate only weakly (|⍴| < 0.4) with age (Supplementary Fig. 1). Owing to specific correlation filters, 36% of the GP-clock model CpGs exhibit a very strong age-correlation (|⍴|>0.8). In turn, PhenoAge and Horvath1 contain no such strongly correlated model CpGs (Supplementary Fig. 1). For Horvarth1 clock this might be a consequence of multi-tissue model training or smaller number of measured CpGs (27K array) or both.

In general, epigenetic clocks such as Horvath1 and GP-clock show high accuracy of age prediction with median absolute errors (MdAE) or mean absolute errors (MAE) around a few years. To test whether clock models can accurately predict chronological age solely based on weakly correlated CpGs with no known association with TFBS, we trained 100 elastic net models on 7,138 blood samples from [22] using 10,000 CpGs exhibiting a weak correlation with age (|⍴|< 0.3) and not overlapping with any TFBS. On average, the models show an MdAE of 3.77 years on an independent healthy validation cohort (n=665, age 18-81 years) (Fig. 1C). This result is comparable to that of some second-generation clocks (MdAE PhenoAge of 3.79 years) and outperforms the first-generation clocks (MdAE Horvath1 of 5.95 years).

Also, we iteratively built a series of elastic-net models by sequentially removing CpGs identified and selected by earlier models as age-predictive from the training feature set for subsequent ones. Although the supposedly “best” age-predictive features were repeatedly removed from the feature set, we observed a markedly slow increase in age prediction errors (Supplementary Fig. 2). Only after approximately fifty rounds of feature removal, i.e., removing 88,007 CpG features (averaging ∼1700 CpGs each iteration), did prediction errors reach the error of the random model.

In summary, our analyses indicate that, given a sufficiently large training cohort, even features that are likely to be biologically less impactful and have limited age-correlated signals can still enable ML to produce reasonable predictions. This finding aligns with other studies that suggest noise or other confounders [19] may contribute to the age prediction accuracy of established epigenetic clocks. This indicates that contemporary clock models may offer limited biological insight because they rely on CpG features less relevant to aging.

### Gene expression changes linked to TFBS-associated CpGs

To further analyze the gene-regulatory roles of TFBS-associated features in the human blood aging data [22], we divided the available CpGs into two groups based on whether they overlapped with experimentally verified TFBS or not (see Methods). Overall, after initial pre-filtering (see Methods), 97,670 CpGs were categorized as part of the TF class, and 132,867 CpGs as part of the non-TF class (Fig. 2A, right). Subsequently, we analyzed the general chromatin annotation of the CpGs in both classes using ChromHMM predictions [24]. As expected, most CpGs in the TF class were located in the promoter region (34%), bivalent promoters (15%), active enhancers (15%), and transcription start sites (TSSs, 11%; Fig. 2A, left). Conversely, CpGs in the non-TF class were found in Polycomb-repressed regions (15%), active enhancers (15%), and bivalent promoters (11%). As a direct consequence of this enrichment, we found that twice as many blood-expressed genes (n = 3380 vs. n = 1698) were linked to CpGs in the TF class compared to the non-TF class (Fig. 2A, right). Using human blood expression data from six age groups spanning 20-80 years in the GTEx database, we next examined how well gene expression in both classes correlates with age. Genes linked to CpGs in the TF class showed a significantly more negative correlation, with a median ⍴ = -0.76 compared to the non-TF class median ⍴ = -0.49 (Wilcoxon test p- value = 2.2e-16, Fig. 2B). Additionally, genes associated with TF class CpGs exhibit higher variability, with a median standard deviation of 0.28 versus 0.03 in the non-TF class (Fig. 2B).

**Figure 2.**
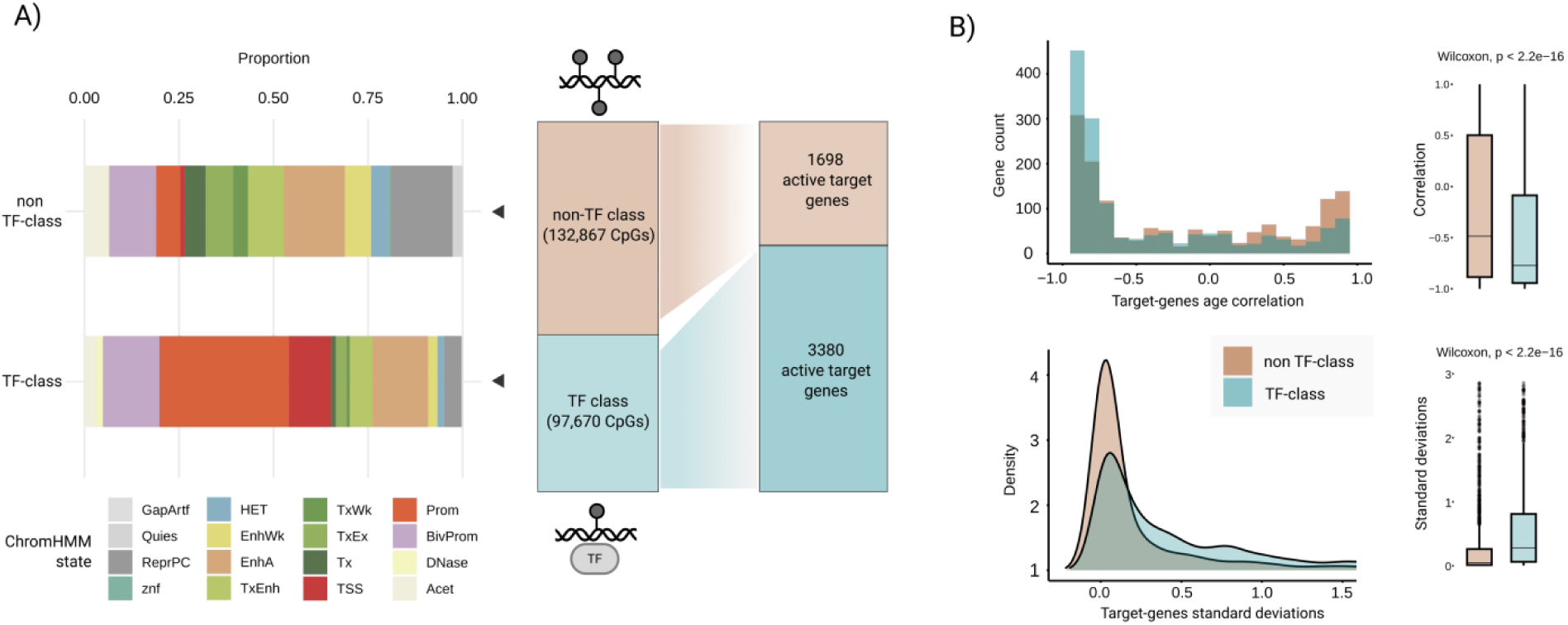
Effects of TF binding at the CpG loci and their target genes during aging. A) CpGs were grouped into two classes based on their overlap with TFBSs, i.e., TF class and non-TF class, respectively. Left: Stacked barplots representing ChromHMM annotations for CpGs in TF class and in non-TF classes. Right: Number of target genes associated with CpGs from TF class and non-TF class that are expressed in blood. B) <u>Top</u>: Histogram (left) and boxplots (right) of Spearman’s correlations of gene expression and chronological age for genes associated with CpGs in the TF class (blue) and the non-TF class (brown). The median Spearmans’ correlation scores of the TF class target genes with age is -0.76 (Q1, 25%: -0.93, Q3, 75%: -0.09) compared to -0.49 (Q1, 25%: -0.89, Q3, 75%: 0.50) of the non-TF class. <u>Bottom</u>: Density plot (left) and boxplots (right) of gene expression deviations during aging for genes associated with CpGs from the TF class (blue) and non-TF class (brown). The median standard-deviation score in the TF class class is 0.28 (Q1, 25%: 0.05, Q3, 75%: 0.81) compared to 0.03 in the non-TF class class (Q1, 25%: 0.001, Q3, 75%: 0.27). Black dots in the boxplots represent outliers (> 1.5 * IQR).

While this result may not be surprising, it highlights that the selection of TFBS-associated CpGs enriches critical regulatory regions, exerting more potent effects on gene expression. More importantly, genes associated with the TF class correlate more strongly with age, exhibit a higher variability, and show a general tendency towards age-related downregulation.

### Age-associated DNA methylation differences at TFBSs

Next, we examined whether age-related DNA methylation changes affect specific TFs more than others. In total, we analyzed 94 distinct human TFs active in blood that bind to CpGs from the TF class. In the first step, using blood data, we calculated the correlation between DNAm changes and age, and identified 14,006 strongly age-correlated CpGs (|⍴| > 0.5). Subsequently, for each TF, we determined the relative frequency of TFBSs that overlap one of these strongly age-correlated CpGs compared to all TFBSs overlapping with any CpG. For comparison, we also calculated the relative frequencies of TFBS overlapping a random set of CpGs of the same size, averaged over a hundred permutations. Among others, TFBSs for ZBED1, IRF8, and NFE2 have a larger proportion of age-correlated CpGs (>20%) compared to random CpGs (∼13%). In contrast, we found that the TFBSs for USF2, CREB1, and TP53 are depleted in age-correlated CpGs compared to the random CpG set (Fig. 3, left). Based on sampling distributions derived from 100 random sets of CpGs from the TF class (Fig. 3, right), we tested for the significance of proportions of age-correlated CpGs in the binding sites of each TF (see methods). In total, we found 14 statistically significant enrichments for TFs (adj. p-value < 0.05), of which eight were enriched for age- associated methylation changes (ZBED1, NFE2, CEBPB, FOXP1, EGR1, SP1, PAX5, and MAZ). In contrast, the binding sites of RBPJ, NFIC, RELA, IKZF1, STAT3, and USF2 were significantly depleted in age-associated CpGs (Fig. 3, right). Interestingly, the expression of ten of the 14 TFs themselves was very strongly correlated with age (|⍴|> 0.9) - increasing for EGR1 and RELA, and decreasing for ZBED1, NFE2, SP1, PAX5, MAZ, IKZF1, STAT3, and USF2.

**Figure 3.**
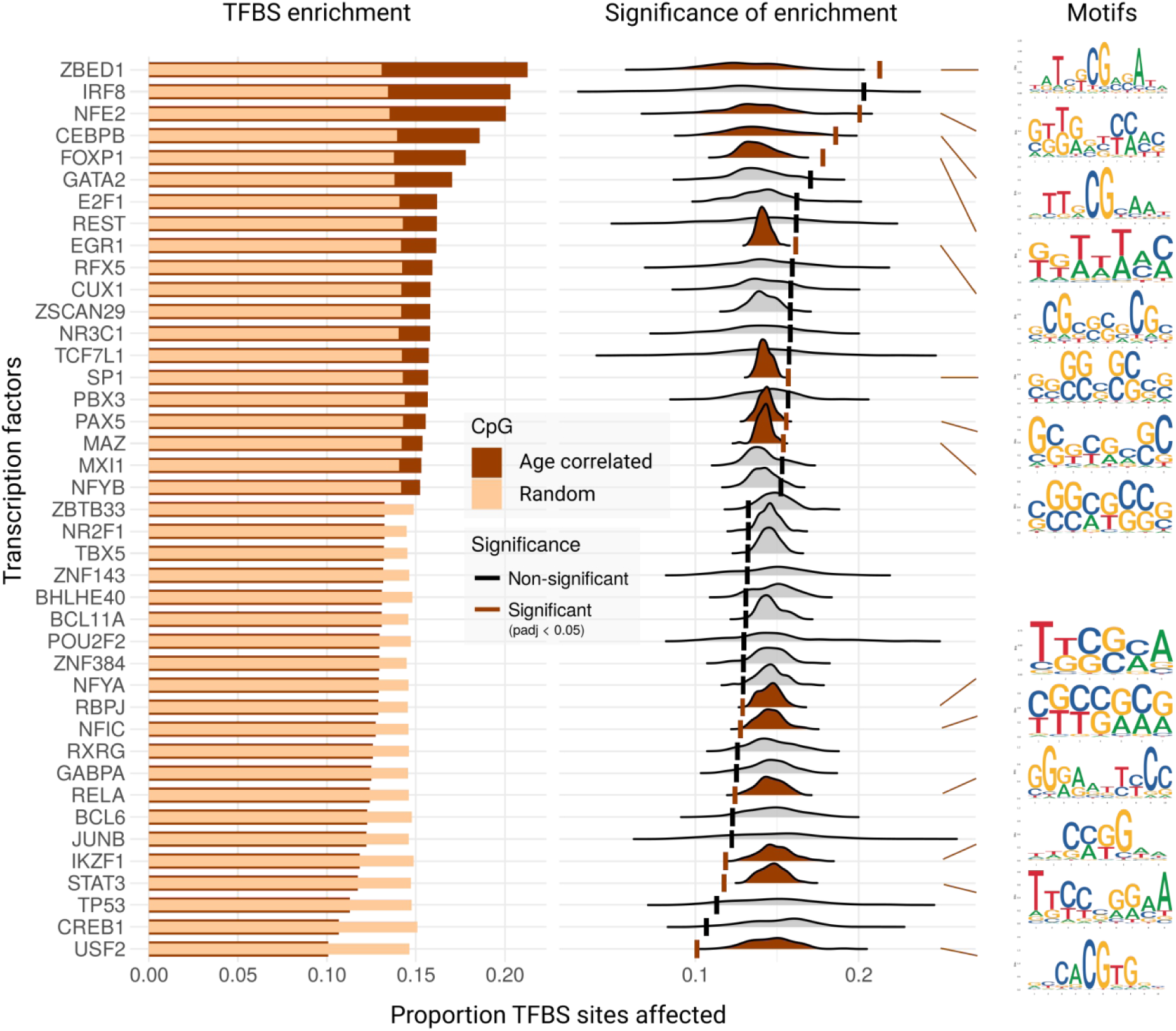
The enrichment of age-associated CpGs in TFBSs. <u>Left</u>: Comparison of TFBS enrichment for age- associated CpGs (with correlation |⍴| > 0.5) in TFBS. Bar plots show the proportion of transcription factor binding sites (TFBSs) affected by age-associated CpGs (dark brown) and by an equal number of randomly sampled CpGs (light brown) across selected TFs. Only the top 20 TFs with the highest observed enrichment and the bottom 20 with the lowest enrichment are shown. The x-axis indicates the proportion of affected TFBSs, and the y axis lists TFs ranked by enrichment. <u>Middle</u>: Enrichment of age-associated CpGs in TFBS. Ridge density plots show the distribution of the proportion of TFBSs affected by randomly sampled non-age-associated CpGs (n=100 iterations) for each transcription factor (TF). Vertical lines represent the observed proportion of TFBSs affected by age- associated CpGs. TFs with significant enrichment (p_adj<0.05) are highlighted in brown. The x-axis indicates the proportion of affected TFBSs, for individual TFs. <u>Right</u>: Age-associated CpG enriched motifs for each TF.

The change in TF gene expression, combined with the methylation changes at its binding sites, indicates that the TFs could govern the expression of downstream genes in an age-dependent manner. Notably, five of these 14 TFs (CEBPB, EGR1, SP1, RELA, and STAT3) are listed by GenAge [25] as human aging-associated genes from experimental studies. In particular, CEBPB, EGR1, and SP1, identified as DNAm-affected in our analysis, play a role in regulating fat metabolism, acute stress response, and general transcriptional control [25]. In contrast, STAT3 and RELA, whose binding sites are depleted in age-associated methylation changes, were functionally linked to chronic inflammation and immune dysregulation, respectively [26];[27]. Upon closer inspection, we found a small fraction of RELA binding sites (12%) undergoing substantial DNAm changes during aging (|⍴| > 0.5). Although not statistically significant, genes in the vicinity of these TFs often showed larger gene expression changes with age as compared to genes with RELA binding sites that exhibited only weak |⍴| < 0.5 DNAm changes (Wilcoxon, p = 0.089, Supplementary Fig. 3). Among the potential RELA targets with substantial DNAm changes, *IRAK4*, *OSM*, and *DUSP2* show strong correlation with age (|⍴| ∼0.9, p-value < 0.05) and are involved in immune-related processes [28];[29];[30]. Similarly, we found target genes of STAT3 with strong age correlation (|⍴| ∼0.9, p-value < 0.05) and TFBS methylation changes. Among them are *ING4*, *MAPKAP1*, and *MAP3K5*, which are involved in inflammation [31];[32]. Interestingly, STAT3 was previously reported to contribute to maintaining a youthful epigenetic state and promoting progenitor-like properties as shown in articular chondrocytes through its regulation of DNA methylation patterns [33].

Taken together, we could identify TFs that are vulnerable to DNAm changes during aging at their binding sites and TFs that may be protected from the DNAm changes at their binding sites. Moreover, genes linked to DNAm-vulnerable TFBSs showed also larger correlation with age.

### Accuracy and robustness of TFBS-based epigenetic clock

Given the interesting properties of age-correlated CpGs overlapping with TFBSs, we next developed an epigenetic clock using these features (Fig. 4A). We initially selected CpGs from the methylation array that overlap with an experimentally validated TFBS (TF class, n = 98,474). Then we filtered those for high age-correlation (|⍴| > 0.5, n = 14,006, Supplementary Fig. 4). To further improve predictions and to reduce noise caused by lifestyle choices, environment, or by other factors (such as probe hybridization, melting temperature, or cellular composition) [7], we removed highly variable CpGs, defined by a standard deviation greater than 10% within the same age group (n=1,451, 0.57% of total CpGs) (Supplementary Fig. 4, with an example CpG). To improve the robustness of any model built from this data, e.g., against missing values and technical noise, we additionally clustered the final set of 14,006 CpGs based on their age-associated methylation profiles using k-means (optimized k = 4,800, see Methods) (Supplementary Fig. 5).

**Figure 4.**
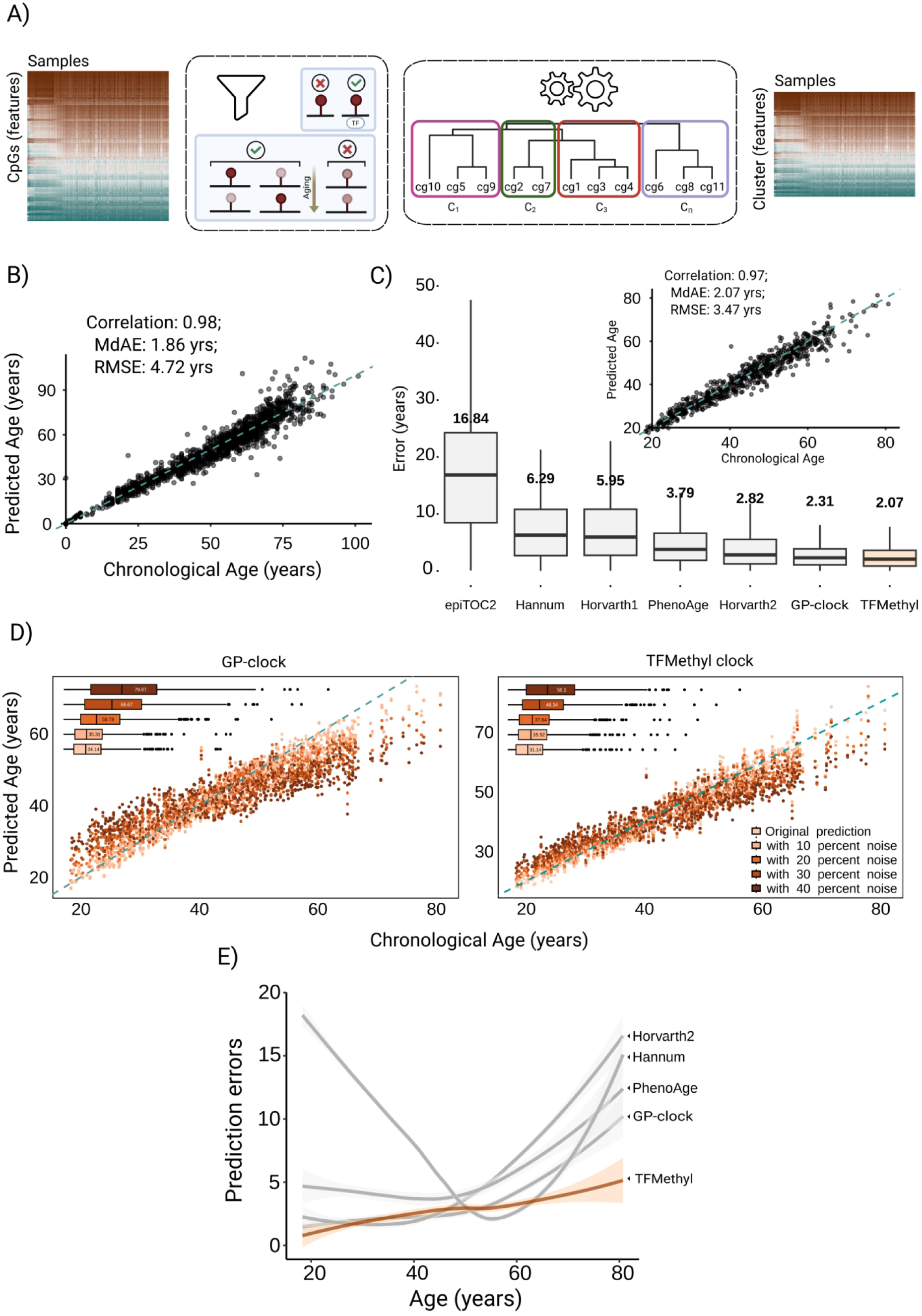
Development and performance of TFMethyl Clock – TFBS-based epigenetic clock model. A) In the TFMethyl Clock architecture, CpGs with large age-correlation and overlapping a TFBS are selected before modelling. Selected features are clustered based on their aging trajectories, and these clusters are used for modeling. B) Predicted age by TFMethyl Clock model (vertical axis) vs the actual chronological age (horizontal axis) on the test set (n = 2143) in years. The dashed blue line represents the perfect positive correlation. C) Distribution of the absolute errors (in years) on the validation cohort (n = 665) for established epigenetic clocks and for TFMethyl Clock, with the median absolute errors (MdAE) above each boxplot. The scatter plot shows the predicted age by TFMethyl Clock vs actual chronological age on the validation cohort. D) Impact of CpG noise on epigenetic age predictions by TFMethyl Clock and GP-clock. Scatter plots show predicted vs. actual chronological ages using the original DNA methylation values (light orange), and using sequentially replacement of 10, 20, 30, and 40% probed CpGs with random values drawn from fitted beta distributions (shades of increasingly darker orange). The dashed blue line represents perfect positive linear correlation. Boxplots depict the median absolute error (MdAE, in months) for each clock under different noise conditions. E) Prediction errors (lines) with 95% confidence intervals (shaded area) by actual chronological age for selected epigenetic clocks (grey) and TFMethyl Clock (light orange).

After completing these preprocessing steps, we trained an elastic net regression model using the median DNAm value of each cluster to predict the chronological age (see Methods). The 4,800 clusters used as input for training had an average cluster size of 3 CpGs. During the training and testing, we employed the DNAm blood Illumina 450k dataset from [22]. The dataset (see Methods, Supplementary Fig. 6), comprising 7,138 samples from 14 studies, was split into a training set (n = 4,997) and a test set (n = 2,141). The model achieved a correlation of r (spearman’s rho) = 0.98 along with comparably low errors (MdAE = 1.86 yrs, RMSE = 4.72 yrs) in the test set (Fig. 4B). Subsequently, we merged the training and test set to train our final age-predictive epigenetic clock model, “TFMethyl Clock”.

This final model, comprising of selected 268 clusters covering 548 CpGs (Supplementary Fig. 7), was evaluated on a held-out validation cohort (GSE84727, n = 665) and compared to established epigenetic clocks. A comparison of our model with the established clocks reveals a pairwise overlap of at most 15 CpGs (3%), with Horvath2 model (Supplementary Fig. 8). In the held-out validation cohort, our model achieved a correlation of 0.97 (MdAE = 2.07 yrs, RMSE = 3.47 yrs) (Fig. 4C). In terms of both the MdAE and RMSE, our model outperforms all other tested clocks. The GP-clock, trained on the same training cohort as ours, was the runner-up with an MdAE of 2.31 years and RMSE of 3.72 years. Horvath2, a clock specifically designed to predict chronological age from blood and skin, achieved an MdAE of 2.82 years. The first-generation clocks Horvath1 (MdAE = 5.95 yrs) and Hannum (MdAE = 6.29 yrs) followed suit. The epiTOC2 clock [34], originally designed to investigate the mitotic age of samples, had the weakest performance on the validation cohort with an MdAE of 16.84 yrs (Fig. 4C).

To test the robustness of our cluster-based model, we randomly selected 10 to 40% of methylation array probes (25,403 to 101,612 CpGs) and replaced their original DNAm values in the validation samples with random values from a Beta distribution fitted to the empirical shape parameters of a particular CpG site (see Methods). We then compared the changes in the performance of the GP-clock and our model on the validation set. The errors of the GP-clock [22] rapidly increased with the addition of noise. In particular, the MdAE increased from 34, 51, 67, and 80 months when replacing 10%, 20%, 30%, and 40% array CpGs, respectively. In turn, our model demonstrated relatively stable performance, with the MdAE increasing by no more than 21 months upon replacing 40% of array CpGs (Fig. 4D). However, this is expected since the goal of the GP-clock was to develop a lean model, that could be measured by targeted PCR.

It has been reported that most DNAm-based epigenetic clocks perform sub-optimally in predicting samples from extremely young and old individuals [35]. Our validation cohort benchmark aligns with this observation. While the largest errors in young samples were observed for Hannum clock, a constant increase in errors for samples of age 60+ was observed for all studied clocks (Hannum, PhenoAge, Horvath2, and Gaussian, Fig. 4E). Compared to the two middle-age quartiles, Horvath2 showed a 2.7-fold increase in error for the oldest quartile, i.e., the top 25% of the oldest individuals. In the same analysis, PhenoAge and GP-clock had a 1.6-fold and 1.9-fold increase in error, respectively. Meanwhile, our approach showed the lowest increase of 1.16-fold.

Based on these benchmarks, our model performs better than established epigenetic clocks in terms of absolute errors, biases in both young and old ages, and robustness to noise.

### Biological aging signatures reflected by TFBS-based epigenetic clock

To assess our framework’s capacity to achieve biological interpretability, we selected CpGs that are stable predictors of epigenetic-age, by training different models using a 10-fold cross-validation (CV) approach. Then, we extracted features from these models with non-zero coefficients in all the CV folds, from here on called age-predictive CpGs (n = 661).

Subsequently, we analyzed the enrichment of age-predictive CpGs at TFBSs of specific TFs. Compared to the background of input CpGs, binding sites of ATF2, EBF1, NR2C2, and REST showed the most substantial enrichment (Fisher’s exact test, p-value <0.05), suggesting a regulatory involvement in age-predictive methylation changes (Supplementary Fig. 9). The corresponding Gene-TF network revealed NR2C2 harboring 221 age-predictive CpGs in its binding sites across the genome. For 41 CpGs, we also found that the expression values of their associated genes strongly correlate with age (|⍴| > 0.8, Fig. 5A).

**Figure 5.**
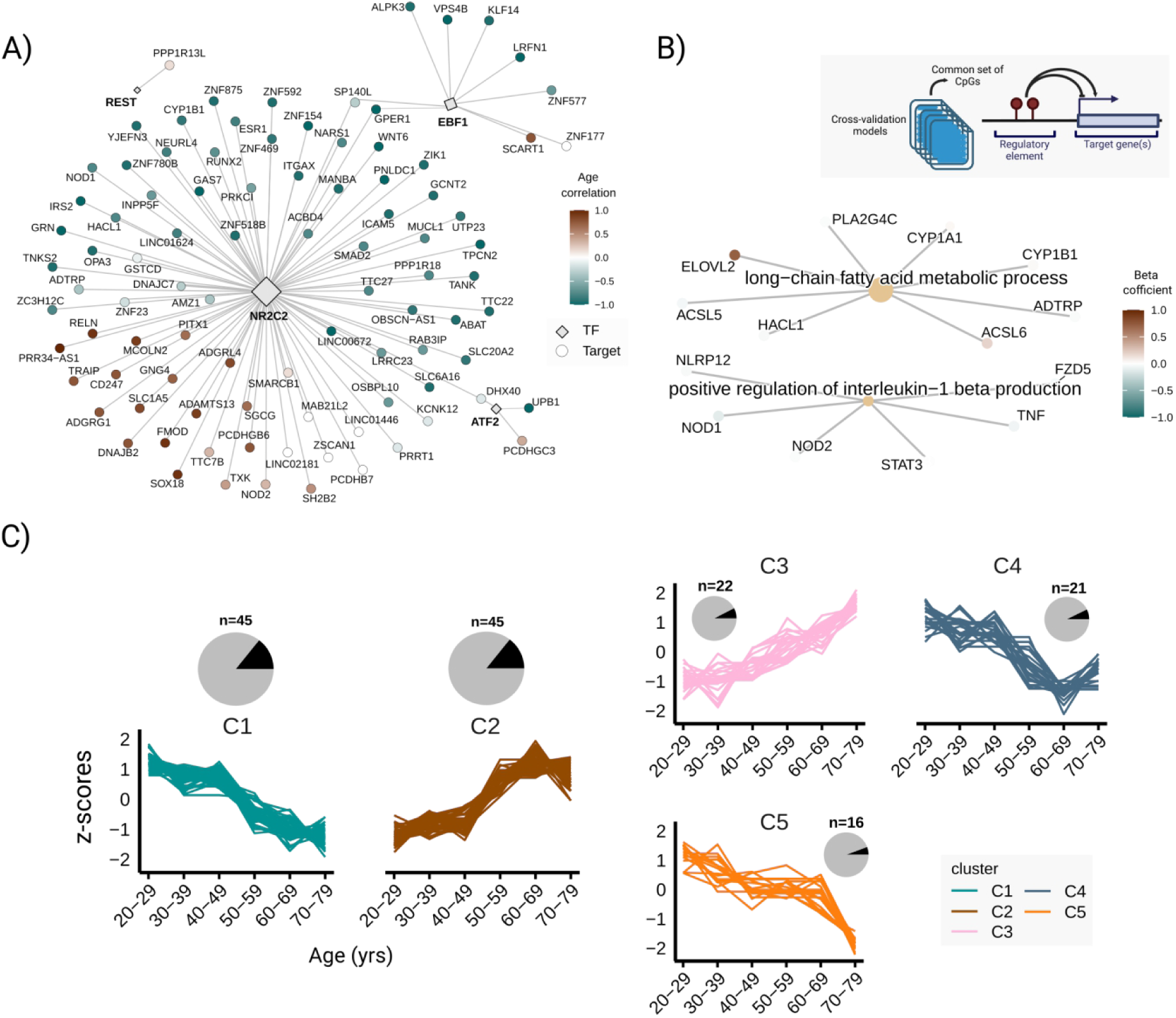
Target genes of age-predictive CpGs selected by TFMethyl Clock. A) Transcription factors are significantly enriched for age-predictive CpGs connected with their target genes. Node size reflects the target-gene set of each TF in the dataset, indicating its regulatory influence. Edges denote directed regulatory relationships from TFs to target genes via corresponding CpGs. Gene node color represents the gene-expression correlation with age. B) Gene Ontology (GO) enrichment analysis for target genes age-predictive CpGs. The size of the GO term node represents the number of genes associated with the respective GO term. The color gradient indicates the direction and magnitude of the averaged model beta coefficients. Only genes TPM > 0.01 were included in the enrichment analysis. C) Gene expression trajectories of target genes of age-predictive CpGs across age groups. Each plot displays the expression dynamics (z-scored) in five distinct co-expression clusters (C1—C5) based on whole-blood RNA-seq data. Pie charts represent the size of each cluster relative to all the genes.

Building on these findings, we analysed all 267 blood-expressed genes that are potentially affected by the 661 age-predictive CpGs (see Methods, detailed in Supplementary table 1). In summary, we found a significant enrichment in pathways (adjusted p-value = 0.02) associated with the positive regulation of interleukin-1 beta production (NOD1, NOD2, STAT3, TNF, FZD5 and NLRP12) and long-chain fatty acid metabolic process (ACSL5, ACSL6, ADTRP, CYP1B1, PLA2G4C, CYP1A1, HACL1, and ELOVL2, Fig. 5B). Strikingly, it has been shown that the ELOVL2, a gene involved in the elongation of polyunsaturated fatty acids, correlates with DNAm changes during aging across various tissues, making it one of the most potent biomarkers for aging [36];[37]. The ELOVL2 gene is annotated to the CpG site cg16867657, which exhibits the largest average coefficient (0.72, nearly double of second biggest cofficient) in our models, indicating a strong age-associated hypermethylation.

To validate the influence of target genes associated with these age-predictive CpGs on the dynamics of the aging process, we examined their expression patterns in the whole blood data from the GTEx collection, comprising 581 samples from individuals aged 20 to 80 years. Notably, the data was obtained from individuals different than the DNAm cohort. Comprehensively, 200 out of all 267 (∼75%) potential target genes had an age-Spearman correlation larger than > 0.5, (Supplementary Fig. 10). Subsequently, we grouped the samples into 10-year age intervals, calculated the median expression value within the group for each gene, and then applied the clust algorithm [38] to obtain gene clusters with similar expression trajectories during aging. After two rounds of clustering, we got five distinct gene clusters (C1-C5), comprising 56% (n=149) of the blood expressed genes in our target set (Fig. 5C). The corresponding methylation trends for these monotonic gene-expression clusters (Supplementary Fig. 11) reveal a somewhat mixed increasing or decreasing trend with age, overall suggesting a locus-specific gene regulation. The largest cluster (C1, n=45) exhibited a linear downregulation during aging and comprised genes such as *IRS2*, *CYP1B1*, and NLRP12. The clusters C2 (n = 45) and C3 (n = 22) showed a non-linear increase in expression during aging and comprised genes such as *PLA2G4C*, *ACSL6*, *NOD2* (C2), and *ACSL5* (C3) (Supplementary Fig. 12).

In particular, C3 contains genes that undergo an expression surge after the age of 60, whereas only a minor increase in expression can be observed before the age of 50. In contrast, cluster C4 (n = 21) exhibits a sudden downregulation between age 40 and 60 following a comparably stable expression in early life (example gene *PPP1R12C* shown in Supplementary Fig. 12). Lastly, the C5 (n = 16) is characterized by stable expression until midlife, followed by rapid downregulation after the age of 60, as is the case for *CYP1A1* (Supplementary Fig. 12). The non-linear trajectories have been recently described in several studies in multi-omics, including DNAm and transcriptome during aging in both humans and mice [39];[40]. Notably, we found several age-predictive CpGs whose target genes also display non-linear expression changes, supporting recent findings on the nonlinearity of aging [39].

## Discussion

Here, we provided evidence that models with CpGs depleted in critical regulatory regions and even lacking strongly correlated aging signals can produce excellent age predictions. This finding draws parallel with previous reports showing that just an increased age-associated noise is sufficient to build accurate aging clocks [41]. Therefore, it may be less surprising that investigations into the biological explainability of DNAm clocks often remained inconclusive [3];[42];[43]. To remedy this shortcoming, we have investigated the potential to build models with CpGs that particularly overlap TFBSs, and have a higher individual correlation with age. This filtering of CpGs limits the feature space and consequently reduces the possible inconsistency of feature selection in elastic net. Interestingly, the applied constraints not only led to improved performance on the test data but also in the independent validation cohort. Among the tested clocks, our TFMethyl Clock model displays a superior performance in terms of chronological age predictions. Instead of training on CpGs directly, we generated CpG meta-features using a simple k-means clustering. This attribute of our approach further alleviates the n>>p problem, deals more effectively with the multicollinearity of CpG features, and, in our view, is more intuitive than other strategies, such as principal component (PC) based strategies, e.g., used in PC clocks [44]. This may contribute to TFMethyl Clock’s robustness to noise and performance for samples at the extremes of the age spectrum, unlike most other epigenetic clocks.

More importantly, our strategy led to the identification of TFBS for specific TFs that significantly enrich age-correlated CpGs, while others appear to be protected. Among the more frequently affected TFBSs are the binding sites for the Zinc Finger BED-Type Containing 1 protein (ZBED1). The protein encoded in the pseudoautosomal region 1 of the X and Y chromosomes acts as a transcription factor for genes involved in cell proliferation. Additionally, it acts as an E3-type small ubiquitin-like modifier (SUMO) ligase. Belonging to a larger family of related proteins that may use the same binding sites, further research is necessary to understand the potential involvement of ZBED1 or other family members in the aging process.

On the other hand, age-associated CpGs were significantly depleted in binding sites of the transcription factor USF2, which is known for its involvement in cellular senescence [45]. Likewise, the TFBSs of CREB1, a methylation-sensitive TF that can bind to unmethylated sites to upregulate LTR repeat transcription [20], were depleted in age-associated CpGs. Further, we observed that all TFs depleted for age-associated CpGs show an enrichment for LTRs among other transposable elements (Supplementary Fig. 13). While the significant depletion of age-associated CpGs in these TFBSs is challenging to interpret, one might speculate that these TFBSs are subject to tighter control of DNA methylation, which actively protects these regions against age-associated dynamics and activation, at least in the respective cell type. We believe that this result encourages follow-up studies to focus on methylation-based properties of individual TFs while aging.

Notably, three quarters of genes associated with age-predictive CpGs found by TFMethyl Clock also exhibit age-related changes in expression. This likely results from our selection process, which effectively enriches regulatory regions. The analysis of genes linked to age-predictive CpGs revealed enrichments in pathways related to interleukin-1, inflammation, and the modulation of immune response – critical pathways impacted by the aging process [46]. Genes such as *NODs*, *STAT3*, and *TNF* are well- established components of innate immunity and inflammaging [47]; [33]; [48]. The second enriched category, "long-chain fatty acid metabolic process," aligns with emerging new literature connecting lipid metabolism to lifespan regulation and cellular senescence [49]. In particular, the recovery of the *ELOVL2* locus, a gene whose methylation is known to be a robust age biomarker, serves as a positive control in support of our modeling approach. Likewise, we found an enrichment of age-predictive CpGs in binding sites of NR2C2 – a factor that has recently been associated with premature aging phenotype [50].

Tracing the expression trajectories of genes associated with age-predictive CpGs, we identified several gene expression clusters that did not enrich any particular biological function. Interestingly, the majority of the gene clusters exhibited non-linear behavior during aging, typically shifting during midlife or in later life. Further studies are necessary to determine whether these changes can be validated and support previously described non-linearities in aging [39].

In this study, we used methylation data from the Illumina 450k array, which only covers a small portion of the approximately 28 million CpGs in the human genome. Using the latest methylation arrays or applying the presented strategy to sequencing-based methylation measurements could offer a more comprehensive view.

In contrast to other approaches, such as the GP-clock, TFMethyl Clock heavily relies on additional biological information, i.e., comprehensive and accurate TFBS data. This dependency naturally limits its application to well-annotated genomes and is prone to bias. As a direct consequence of its design, it does not consider other regulatory CpGs that are not directly involved in TF binding. Since TF binding is often tissue-specific, we believe studying other tissues is necessary to understand better global, recurrent aging signatures, as well as those that are more tissue-specific. This approach would be especially helpful in understanding the role of the identified transcription factors’ dynamics in aging.

## Methods

### DNAm dataset

The methylation dataset used to build the final predictor combined 15 publicly available DNA methylation cohorts, borrowed as a curated set from [22]. The 7803 samples used were mostly classified as healthy and the cohort was overall sex balanced. The minimum and maximum age of the cohort was 0 and 101 years respectively, while the mean was 39.10 years and median was 40 years. The GSE105018 contributes the maximum 1651 samples to the cohort, and the GSE55763 contributes a minimum of 12 samples (Supplementary Fig. 6). To evaluate the performance on the 30% test split—we divided the entire cohort with training samples n = 4995 after removing a test-cohort of n = 2143 with a similar age distribution, and an untouched validation GSE84727 (n = 665) cohort—similarly as done by [22]. Overall no imputation method was used for calculating the missing CpG probe values, as only the measured probes in all the samples were considered for the further analysis. We also discard the samples where chronological age was unknown or if the sample had missing values for the given common set of probes.

### RNA-seq dataset

The RNA-seq gene-expression data used in the study originates from the GTEx Analysis V10 release cohort, downloaded on 29/11/2024. Whole blood tissue filter was used before downloading the TPM and read count data from the GTEx portal, totalling 581 samples. The age groups analyzed were from 20 to 80 years old, with a median at 55 years. We took the middle of the age-group as the respective age for each sample (for example, 25 for 20-29 yrs). Atleast 35% of the cohort here was reasonably healthy as cause of death before sample collection being sudden or unexpected—largely by accidents or trauma/suicide. Sequencing reads were built on the GRCh38/hg38 genome, STAR aligner (v2.7.10a), and the RSEM (v1.3.3) was used for the transcript quantification by the standard GTEx processing pipeline.

### Transcription factor binding sites (TFBS) annotation

We curated a genome-wide TFBS annotation for all the TFs whose overlapping motifs and binding site information was present in the JASPAR and cistrome databases, respectively [51];[52]. Entire hg38 genome JASPAR TF-motifs and cistrome TF Chromatin Immunoprecipitation-sequencing (ChIP- seq) datasets were downloaded from the respective online repositories. Then for the ChIP-seq TF binding profile, we filtered for “Blood” tissue-type, and further concatenated different ChIP experiments together for each TF using bedtools (v2.30.0-56) [53]. Finally, we intersected ChIP profiles with the motif profiles for each TF using bedtools, thus creating a set of TFBS for all possible TFs. From the total of 828 TF motifs available in JASPAR database and 239 TFs with ChIP-seq binding profiles in blood samples, a final common TFBS set of 94 TFs was selected.

### CpG feature pre-filtering and clustering modules

The initial methylation matrix consists of 254,028 CpGs before any filtering or clustering steps, combining all measured probes in 7803 samples. The first filter for 94 TFs binding sites was intersected in R using GenomicRanges::findOverlaps() function, obtaining 98,474 CpGs. The genomic positions for CpGs were determined using the IlluminaHumanMethylationEPICv2anno.20a1.hg38 R package (v1.0.0) - originally curated from the Illumina EPIC v2 methylation array annotation. The high-variable CpGs (n=1451) were identified and removed from the set using sd from {stats} in R. Further, Spearmans’ correlation coefficients for CpGs with age was calculated using the base R cor{stat} function. Then all the CpGs with cutoff of |r| > 0.5 were retained, resulting in 14,006 CpGs for the clustering algorithm.

These 14,006 CpGs were then grouped into 4800 clusters, forming meta-features for the regression model. The k-means clustering was performed using kmeans {stats} function from the base R. For each of these 4800 clusters, the median centroid methylation value was taken as the feature input to the elastic net-regression algorithm.

### Enrichment analysis of age-associated CpGs in TFBS

Non-overlapping TFBS counts were calculated per transcription factor using GenomicRanges, and observed enrichments for age-correlated CpGs were compared to null distributions generated from 100 random CpG sets of equal size. Normality was assessed with the Shapiro–Wilk test, followed by either a parametric z-test or a non-parametric empirical test, with multiple testing correction by the Benjamini–Hochberg method.

### The prediction model

The algorithm used was from the gaussian elastic-net regression family with 90% ridge (L2) and 10% lasso (L1) penalties, and 30-folds of inner cross-validation to tune the hyperparameter (lambda). The function used to train the elastic-net linear regression model glmnet::cv.glmnet() on the methylation data was from the glmnet R library (v4.1-8) [54];[55]. Importantly, the target variable (chronological-age) was log-transformed to account for exponential-like dynamics (homoscedasticity) for many CpGs methylation with age and obtain a more bell-shaped sample distribution, as shown before [56].

### Contemporary epigenetic clocks

We used R methylCIPHER package (v0.2.0) to run previously published epigenetic age predictors with no imputation setting (“imputation = F”). Specifically, before analyzing age outcome results from Epigenetic Time of Cancer 2 (EpiTOC2) clock, we divided the observations by a hundred to correct for the offset bias. For most of the epigenetic clocks tested, almost all the probes were present in the input methylation dataset, for almost all the samples.

### Beta methylation noise-simulation

To simulate measurement noise, CpG values were perturbed by sampling from a Beta distribution parameterized using the empirical mean (μ) and variance (var). The Beta distribution parameters alpha (α) and beta (b) were estimated as:

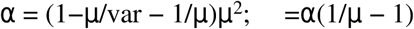

Using these parameters, new methylation values were generated for CpGs using rbeta() function from base R {stats}: xi = rbeta(α,b), i=1,…,665; where 665 corresponds to the number of samples in the validation dataset.

### 10-fold cross-validation setup and target gene-connections

To narrow down to the most important age-predictor CpGs, we performed ten repetitions of stratified leave-one-set-out cross validation. The CV folds were divided using createFolds() function from the caret R library (v7.0-1) having similar age distribution.

We connected the CpGs common between CV models to their associated gene targets through IlluminaHumanMethylationEPICv2anno.20a1.hg38 annotations. We adapted a customized function to associate a single gene target to each CpG, in the case where multiple genes connect to the CpG locus. In total, filtering for genes expressed in blood (average TPM in the GTEx cohort > 0.01), resulted in 267 genes.

### GO enrichment analysis for target genes

The CpG associated gene targets were analyzed for the possible gene ontology enrichment through enrichGO() function from the R library clusterProfiler (v4.12.0). The annotation used for performing the analysis comes from the bioconductor annotation data package org.Hs.eg.db (R library v3.19.1) for the human hg38 annotation. Appropriate enrichment background was established by taking the intersection of human blood expressed genes (average TPM > 0.01), and 450k array CpG associated genes. The enrichment q-value threshold was 0.05 and we used the Benjamini—Hochberg method for p- value adjustment.

### Clustering the target genes using age expression profiles

For the gene targets, we analyzed their expression trajectories during aging by clustering them in different clusters using the clust tool (v1.18.0). All the target genes were clustered based on the Z-scores of the age group median expression with two iterations. After the first clustering round, ∼66% of CpGs associated genes remained unclustered. Hence, another round of clustering was performed resulting in a total of 5 clusters comprising 56% of all genes considered.

### TE distribution analysis within TFBS

To analyze the distribution of transposable elements (TEs) across affected and protected TFs, we used the DeepTools suite. We first applied the computeMatrix function (v3.5.5) in scale-regions mode to generate a matrix of signal intensities surrounding regions of interest. Specifically, we used bigWig files representing transposable elements (TEs) signal tracks for four major TE families: LINE, SINE, LTR, and DNA transposons from the human genome hg38. These tracks were processed against two sets of genomic regions—TFs affected and protected for age methylation changes—provided as BED files. We included 2 kb upstream and 2 kb downstream flanking regions (-a 2000 -b 2000) for each TFBS, and treated any missing data as zero to avoid bias (--missingDataAsZero). The average methylation signal across regions was plotted as a summary track above the heatmap using the mean value.

### Visualization

Plots were created using the following R packages: ggplot2 v3.5.1, ggrastr v1.0.2, ggridges v0.5.6, and ComplexHeatmap v2.20.0. We also used plotHeatmap 3.5.5 to plot the TE density within TFs.

## Supplementary text

While DNA methylation is frequently associated with the silencing of transposable elements (TEs) [20,57], several studies have also indicated that TEs may serve as regulatory hubs harboring a plethora of TFBSs [58]. To investigate the age-related dynamics in this specific context, we overlapped TEs with TFs that are enriched and depleted in age-correlated CpGs, respectively. While LINE and SINE elements were depleted for both age-associated of TFs, long terminal repeats (LTRs) enriched TFBSs with age-resilient CpGs (Supplementary Fig. 13). A possible explanation of this could be that TFBS coincide with LTRs are maintained at higher levels of methylation to keep them transcriptionally silenced by genomic mechanisms, otherwise leads to catastrophic repeat activation [20].

The average model coefficients for the 303 age-predictive CpGs ranged from +0.72 ≤ b ≤ -0.55, with a median value of -0.01 (Supplementary Fig. 10). The most negative coefficient (-0.55) was found for the singleton cluster associated with the CpG cg22982767. The locus is associated with the PRR34- AS1 gene. The most positive coefficient was obtained for cg16867657 associated with the ELOVL2. Eighty-nine of these even had a statistically significant correlation (adjusted p-value < 0.05, 33.3%)— suggesting a potential role of selected CpGs in gene transcriptional regulation during aging.

## Supplementary figures

**Supplementary Figure. 1.**
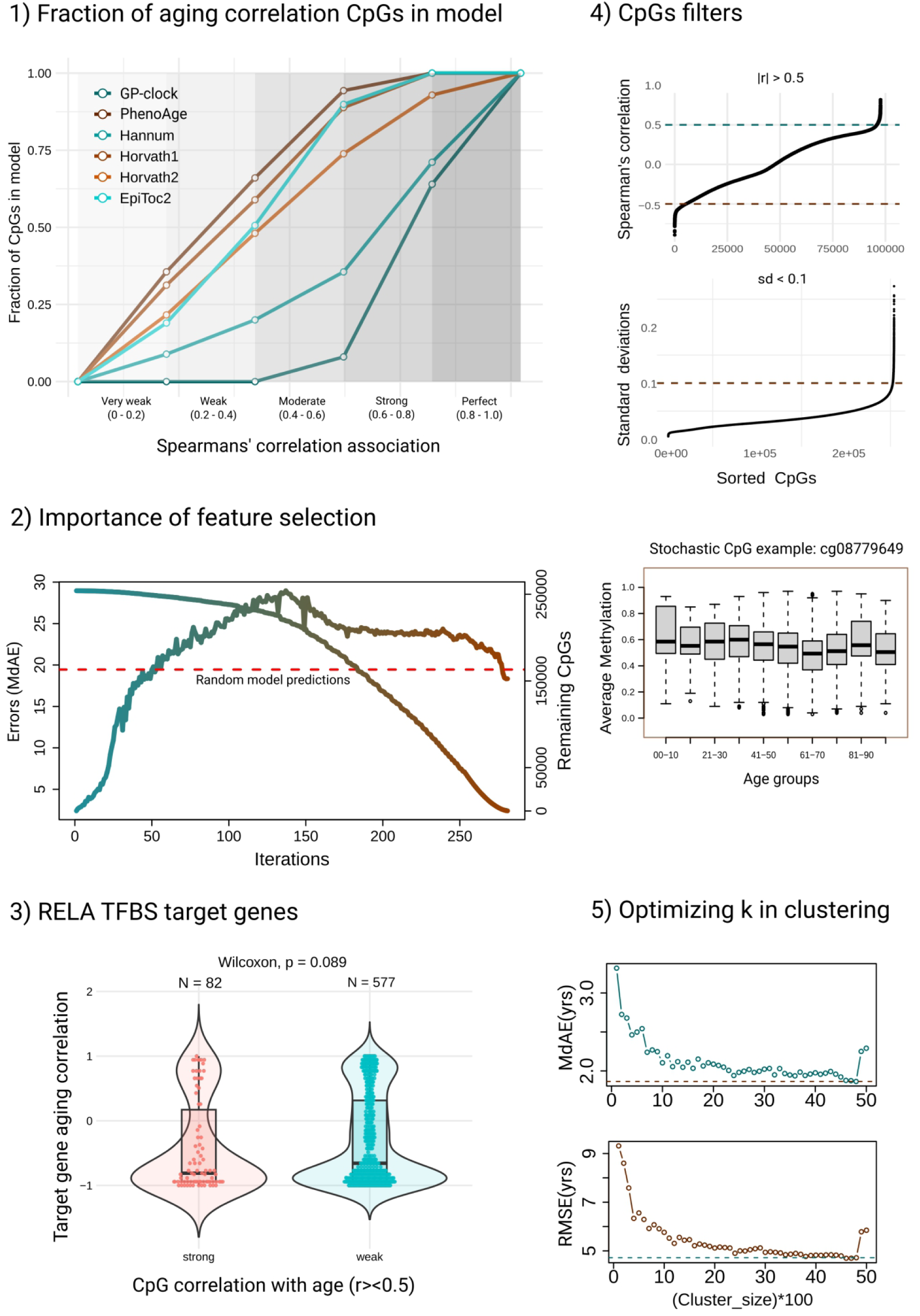
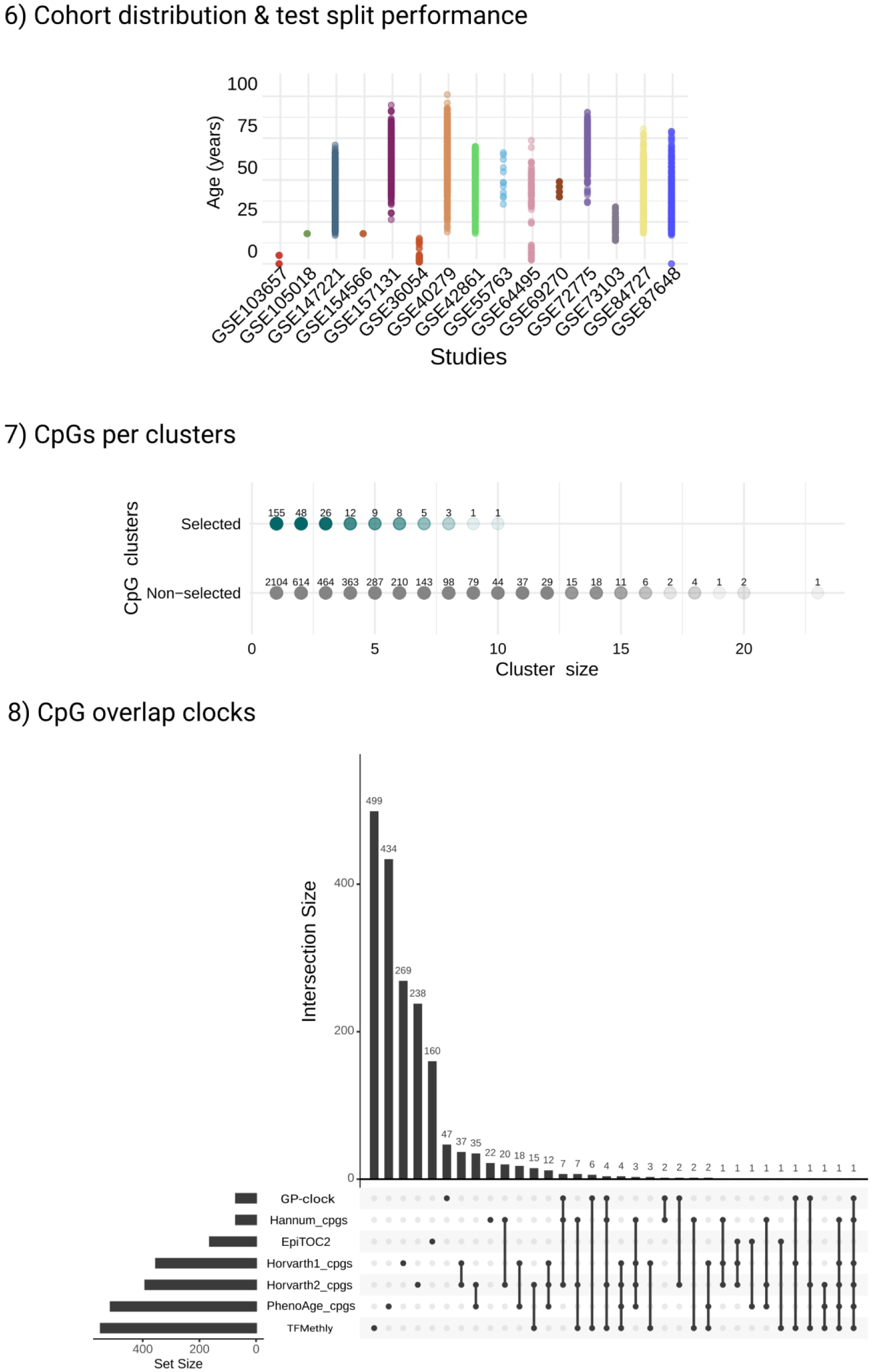
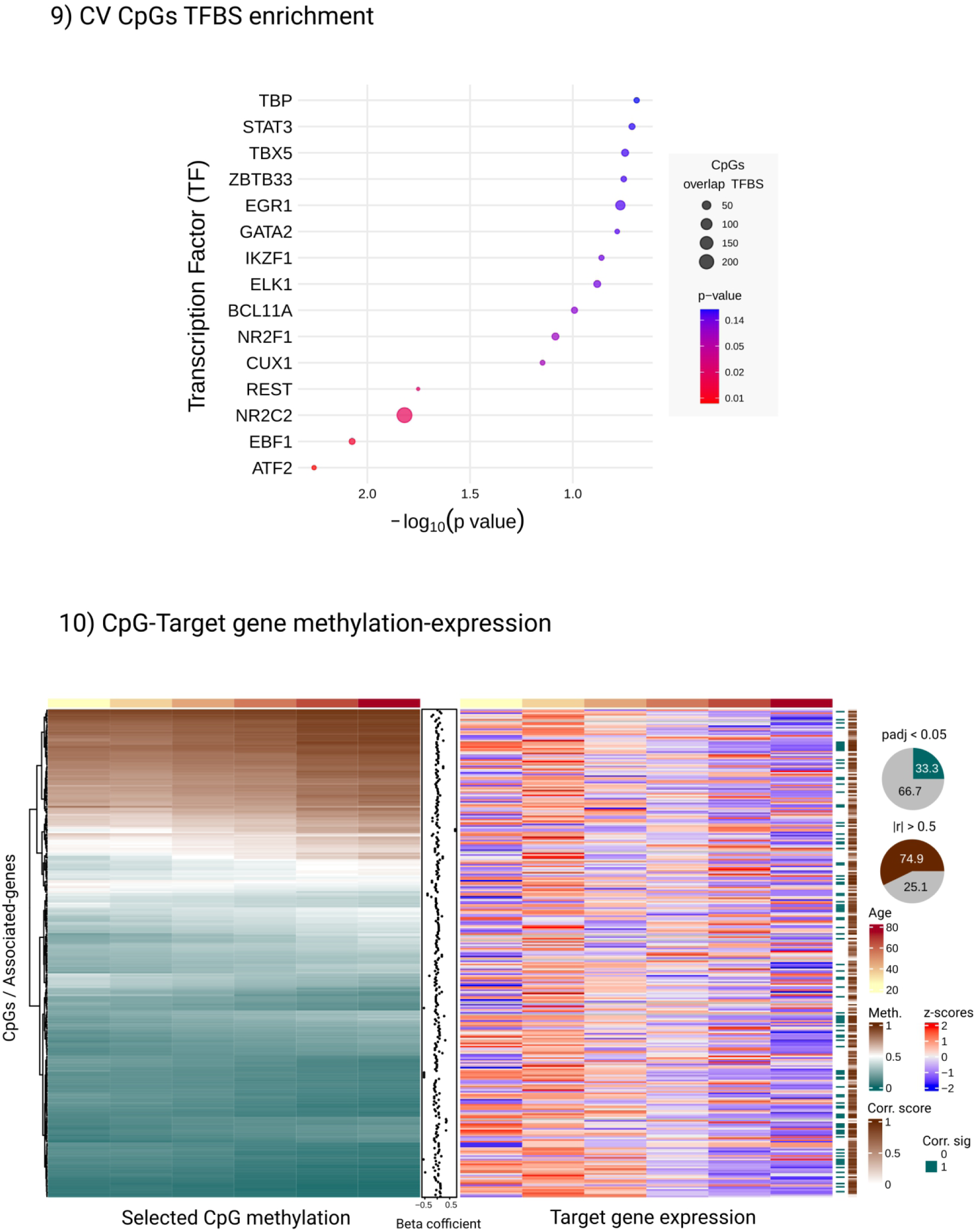
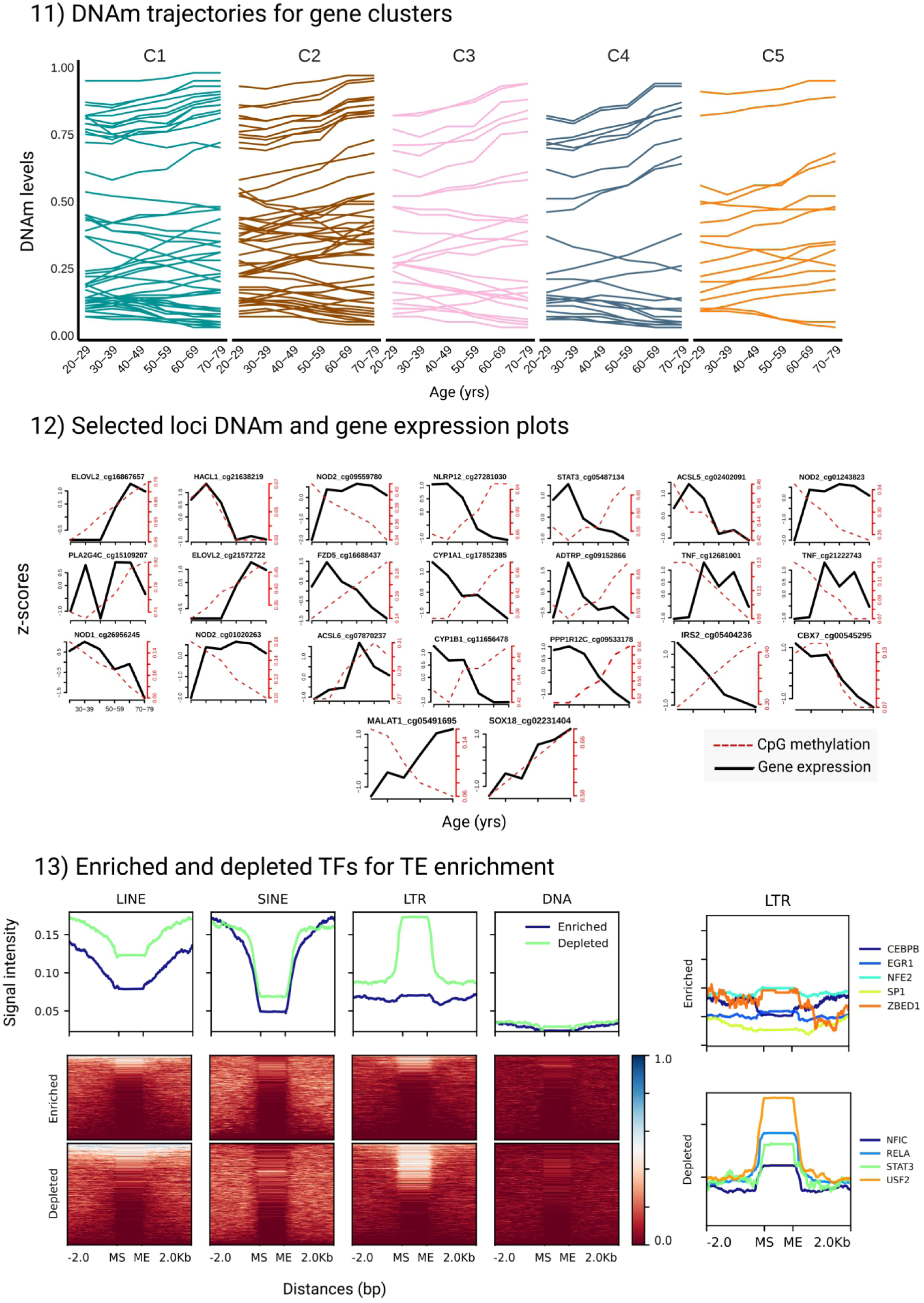
CpG overlaps across aging clocks at different correlation thresholds. Proportion of CpGs from each epigenetic clock retained below increasing correlation cutoffs. Shaded regions denote different correlation intervals. Lines represent six clocks: GP-clock, Hannum, Horvath1, Horvath2, and PhenoAge. 2) Iterative elimination of age-predictive CpGs while training epigenetic age predictors. The plot shows the median absolute error (MdAE) across iterations of elastic net regression models trained on progressively reduced CpG sets. In each round, CpGs with non-zero model coefficients were removed, refining the feature space. The dashed red line indicates the baseline error when predicting all samples as the cohort average, i.e random model. The overlay (right axis) shows the cumulative number of CpGs eliminated over iterations, highlighting the model’s reliance on a shrinking subset of predictive features. 3) Box-violin plot for gene expression correlations of RELA TFBS target genes categorized based on associated CpG methylation change with age. Left is stronger change with age (|r| > 0.5) and right is weaker correlations with age (|r| < 0.5). 4) <u>First</u> (from top): The correlation filter (|r| > 0.5) for CpGs demonstration. Every CpG with spearman’s correlation value above red and blue dotted was further selected. <u>Second</u>: The stochasticity filter for CpGs. Everything above 0.1 red dotted line was removed from the further analysis. <u>Third</u>: Boxplots for an example CpG which was removed. Boxes represent the mean and 95% confidence interval for methylation level at the given loci for a particular age group. 5) Performance of age prediction across cluster sizes. Median absolute error (MdAE) and root mean square error (RMSE) are plotted against increasing cluster sizes used in optimization. Dashed lines indicate performance at the selected optimal cluster size. 6) Cohort age distribution for different GSE studies considered in this study. The vertical axis represents the age for the particular sample from each study on the horizontal axis. 7) Distribution of CpG cluster sizes selected versus not selected by the elastic net model. Each point represents a CpG cluster, colored by whether it was selected by the elastic net model (non-zero coefficients) or not. 8) Overlap of CpGs across multiple epigenetic clocks. UpSet plot showing shared and unique CpGs between various aging clocks, including PhenoAge, Hannum, Horvath1, Horvath2, EpiTOC2, GP-clock, and the newly derived TFMethyl Clock model. Bars represent the number of CpGs present in each intersection set. 9) Top 15 transcription factors enriched among CpG sites consistently selected across 10-fold CV in the aging methylation model. Enrichment was assessed using Fisher’s exact test, with point size indicating the number of overlapping CpGs and color representing the p-value on a log scale. 10) <u>Left</u>: DNA methylation levels at selected CpG sites. Heatmap representing methylation beta values of CpG sites commonly selected across all 10 folds of CV. CpGs were ordered by average model coefficient (weights), shown as a row annotation. Columns represent individual samples, and color intensity indicates methylation level (teal to brown). A top annotation bar shows the chronological age of individuals. <u>Right</u>: Expression of genes associated with selected CpG sites. Heatmap of median gene expression (TPM) per age bin from GTEx whole blood RNA-seq data, converted to z-score normalized across samples. Genes were annotated to CpG sites via the Illumina EPIC v2 annotation and ordered to match the CpG heatmap. Right-side annotations include: Corr. sig as the adjusted significance of Spearman correlation with age (FDR < 0.05, teal); Corr. val as absolute Spearman correlation coefficient |ρ|, scaled from white to brown. Pie chart showing the percentage of genes with FDR-adjusted p-value < 0.05 from Spearman correlation tests with age (teal) vs. non-significant genes (grey). Pie chart indicating genes with absolute Spearman correlation coefficient greater than 0.5 (brown) versus all others (grey). 11) Line plots show methylation level dynamics of CpG sites associated with gene clusters C1—C5 across five age bins: 20—29, 30—39, 40—49, 50—59, 60—69, and 70—79 years. 12) The line plots representing selected loci gene-expression and DNAm change within age-groups from young to old, left to right on the x-axis. The y1 left axis denotes z-scores for gene expression, the y2 axis denotes DNAm level from increasing methylation (bottom to top). 13) Transposable element signal enrichment around affected and protected TFs. Heatmaps and average signal profiles display the distribution of four major transposable element (TE) classes—LINE, SINE, LTR, and DNA transposons—around TFs either enriched or depleted for methylation-associated signals (±2 kb from motif start or end). Signals were extracted from corresponding bigWig files using computeMatrix scale-regions, with missing values imputed as zero. The color scale reflects normalized signal intensities (red = high, blue = low), and the average profile above the heatmap shows the mean signal across all regions for each TF class. MS: motif start; ME: motif end.

